# Can Artificial Intelligence Detect Monkeypox from Digital Skin Images?

**DOI:** 10.1101/2022.08.08.503193

**Authors:** Towhidul Islam, M.A. Hussain, Forhad Uddin Hasan Chowdhury, B.M. Riazul Islam

## Abstract

An outbreak of Monkeypox has been reported in 75 countries so far, and it is spreading at a fast pace around the world. The clinical attributes of Monkeypox resemble those of Smallpox, while skin lesions and rashes of Monkeypox often resemble those of other poxes, for example, Chickenpox and Cowpox. These similarities make Monkeypox detection challenging for healthcare professionals by examining the visual appearance of lesions and rashes. Additionally, there is a knowledge gap among healthcare professionals due to the rarity of Monkeypox before the current outbreak. Motivated by the success of artificial intelligence (AI) in COVID-19 detection, the scientific community has shown an increasing interest in using AI in Monkeypox detection from digital skin images. However, the lack of Monkeypox skin image data has been the bottleneck of using AI in Monkeypox detection. Therefore, in this paper, we used a web-scrapping-based Monkeypox, Chickenpox, Smallpox, Cowpox, Measles, and healthy skin image dataset to study the feasibility of using state-of-the-art AI deep models on skin images for Monkeypox detection. Our study found that deep AI models have great potential in the detection of Monkeypox from digital skin images (precision of 85%). However, achieving a more robust detection power requires larger training samples to train those deep models.

## I. Introduction

MONKEYPOX virus has been spreading throughout the world at a rapid rate, while the world is still recovering from the aftermath of Coronavirus disease (COVID-19). Monkeypox is an infectious disease and its recent outbreak has been reported in at least 75 countries to date. The first human infection by the Monkeypox virus was reported in the Democratic Republic of Congo (formerly Zaire) in 1970 [1]. The Monkeypox virus belongs to the genus *Orthopoxvirus* of the family *Poxviridae* [2], which was first transmitted from animals to humans. Typically, Monkeypox infection shows symptoms similar to those of Smallpox infection [1]. Smallpox was largely eradicated in 1970, leading to the cessation of Smallpox vaccination. Since the 1970s, Monkeypox has been considered the most dangerous *Orthopoxvirus* for human health. In the past, Monkeypox was most frequent in the African continent, however, it has often been reported in urban areas outside the African continent now-a-days [1]. Scientists believe that the current Monkeypox outbreak in humans on a global scale is due to changes in biological attributes of the Monkeypox virus, changes in human lifestyle, or both [3].

The clinical attributes of Monkeypox are similar to that of Smallpox, however less severe [4]. On the other hand, skin lesions and rashes, caused by Monkeypox infection, often resemble those of Chickenpox and Cowpox. This clinical and visual similarity among different pox infections makes the early diagnosis of Monkeypox challenging for the healthcare professional. In addition, the rarity of Monkeypox infection in humans before the current outbreak [5] created a knowledge gap among healthcare professionals around the world. For the diagnosis of Monkeypox infection, the polymerase chain reaction (PCR) test is generally considered the most accurate tool [6]. However, healthcare professionals are often used to diagnosing pox infections by visual observation of skin rashes and lesions. Monkeypox infection has a low mortality rate (i.e., 1%-10%) [7], however, early detection of Monkeypox may help in patient isolation and contact tracing for effective containment of community spread of Monkeypox.

Different artificial intelligence (AI) tools, especially deep learning approaches, have been widely used in different medical image analysis tasks (e.g., organ localization [8], [9], organ abnormality detection [9], [10], gene mutation detection [11], cancer grading [12], [13] and staging [14]) in the last decade. Notably, AI methods have recently played a significant role in COVID-19 diagnosis and severity ranking from multimodal medical images (e.g., computed tomography (CT), chest X-ray, and chest ultrasound) [15]–[17]. This success motivates the scientific community in utilizing AI approaches for Monkeypox diagnosis from the digital skin images of patients. It is well-known that supervised or semi-supervised AI approaches are data-driven, which require a large number of data for developing AI methods effectively. However, there is no publicly available and reliable digital image database of Monkeypox skin lesions or rashes. To urgently tackle this situation, we used the web scraping approach (i.e., extracting data from websites) [18] approach to collect digital skin lesion images of Monkeypox, Chickenpox, Smallpox, Cowpox, and Measles to facilitate the development of AI-based Monkeypox infection detection algorithms. However, the immediate question follows if we can utilize state-of-the-art AI techniques on digital skin images of rashes and lesions to accurately identify Monkeypox and distinguish it from infections by other types of poxes and diseases (e.g., Chickenpox, Smallpox, Cowpox, Measles, etc.). To our knowledge, only two AI-based Monkeypox detection studies have been conducted to date that appear as preprints [19], [20]. However, these studies have several limitations. First, these studies included only three cases of disease (i.e., Monkeypox, Chickenpox, and Measles). Second, these studies are conducted on very small datasets. Third, these studies tested one [19] or a few [20] AI deep models in pox classification tasks.

In this paper, we test the feasibility of using state-of-the-art AI techniques to classify different types of pox from digital skin images of pox lesions and rashes. The novelties of this work are the following.

1. We utilize a database that contains skin lesion/rash images of 5 different diseases (compared to 3 diseases in [19], [20], i.e., Monkeypox, Chickenpox, Smallpox, Cowpox, and Measles, as well as contains healthy skin images.
2. Our database contains more data for pox, measles, and healthy images scraped on the Web (i.e., 804 images), before augmentation, compared to other similar databases used in [19], [20] (204 and 228 images, respectively).
3. We tested the disease classification power of seven state-of-the-art deep models (compared to one and three deep models in [19] and [20], respectively) from digital skin images. We tested the disease classification performance of ResNet50 [21], DenseNet121 [22], Inception-V3 [23], SqueezeNet [24], MnasNet-A1 [25], MobileNet-V2 [26], and ShuffleNet-V2 [27].
4. We performed 5-fold cross-validation tests for each of the AI deep models to more comprehensively analyzing our findings, which is not done in previous studies [19], [20], although their dataset is smaller than ours.

## II. Methodology

In this section, we describe our approach to data collection, data augmentation, and experimental setup.

### A. Data Collection

#### 1) Web-scraping for Image Collection

We use web scraping to collect skin infected with Monkeypox, Chickenpox, Smallpox, Cowpox, and Measles, as well as healthy skin images from various sources, such as websites, news portals, blogs, and image portals using the Google search engine. We show the pipeline of our database development in Fig. 1. We searched to collect images that fall under “Creative Commons licenses.” However, for several pox classes, we hardly find images. Therefore, we also collect images that fall under “Commercial & other licenses.” Therefore, we include supplementary material that includes the uniform resource locator (URL) of the source, the access date, and the photo credit (if any) for all our collected images. In Fig. 2, we show some example images from our database.

**Fig. 1.**
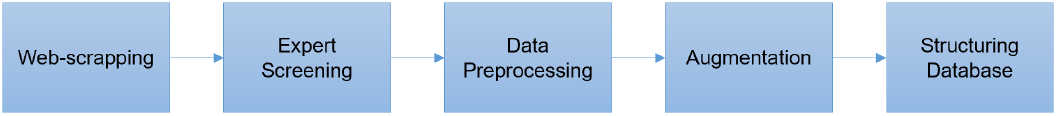
The pipeline of our database development.

**Fig. 2.**
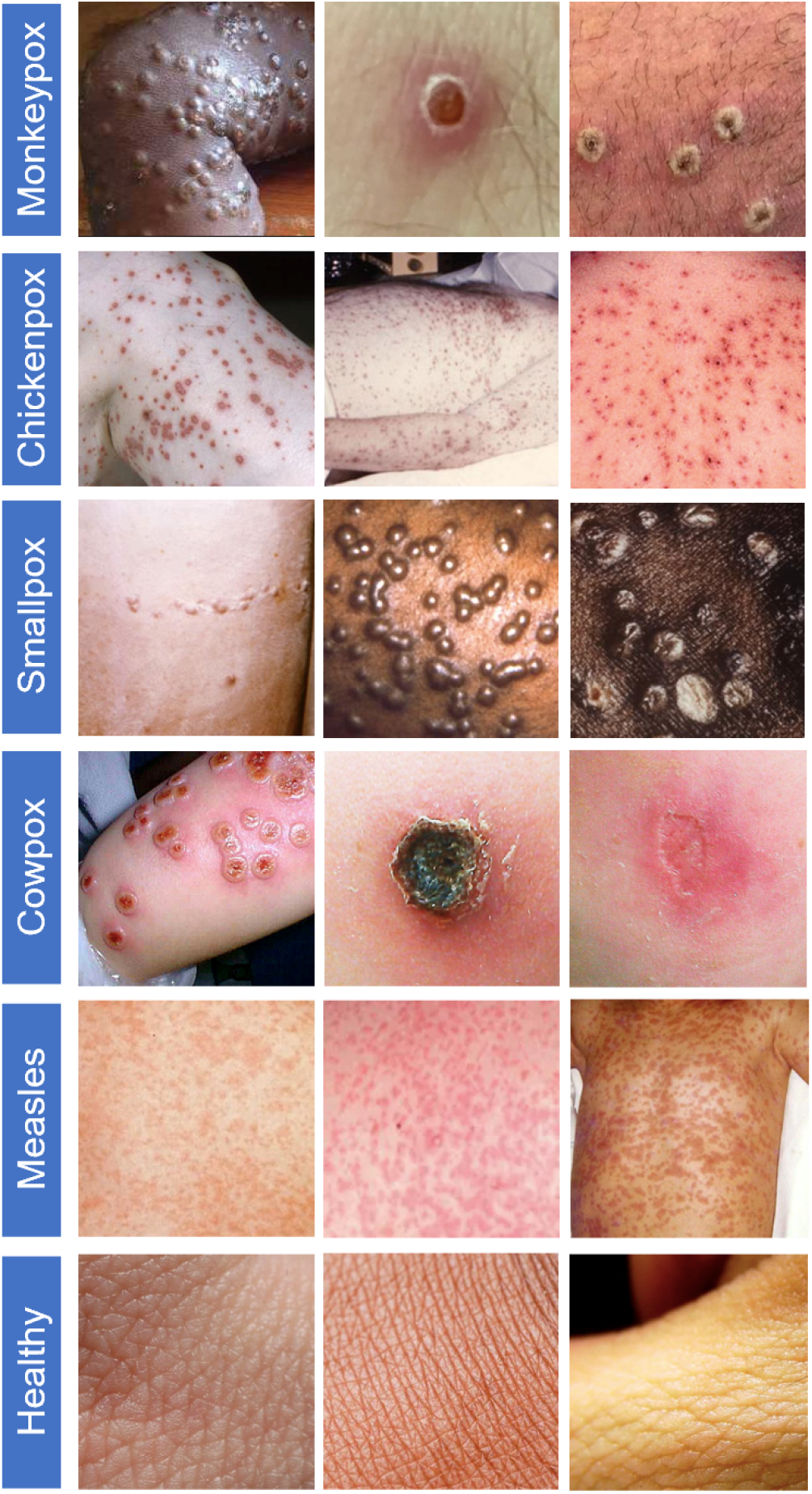
Example skin images of Monkeypox, Chickenpox, Smallpox, Cowpox, Measles, and healthy cases (first to sixth rows, respectively) from our database.

#### 2) Expert Screening

Two expert physicians, who were experts in infectious diseases, screened all images collected to validate the supposed infection. In Fig. 3, we show a pie chart of the percentage of original web-scraped images per class in our dataset (after expert screening).

**Fig. 3.**
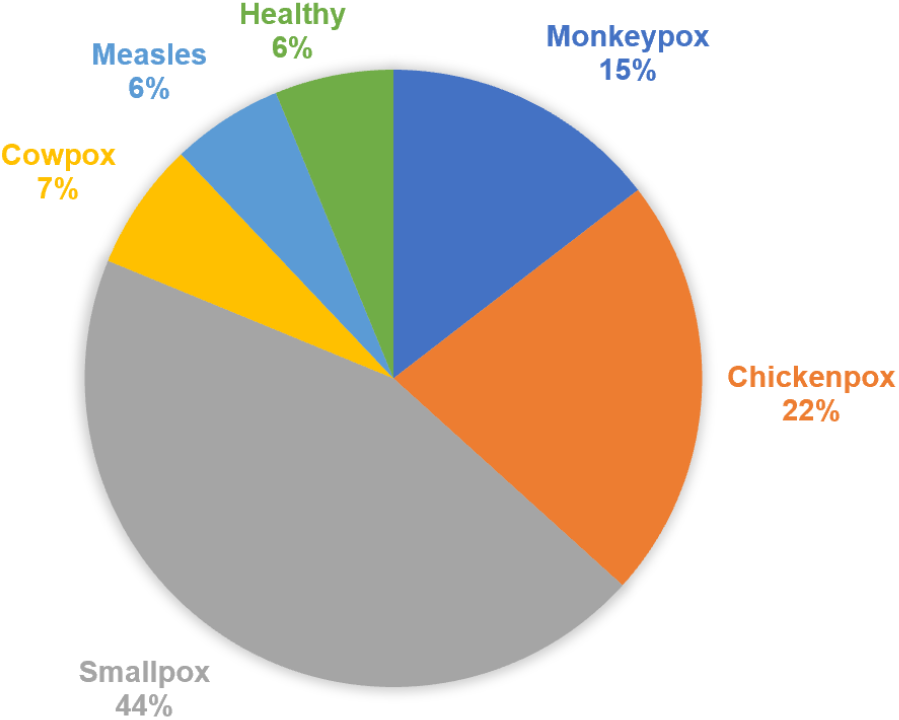
A pie chart showing the percentage of original web-scraped images per class in our dataset.

#### 3) Data Preprocessing

We cropped images to remove unwanted background regions and blocked the eye region with black boxes to make patients nonidentifiable from their corresponding images. We also did the same to hide revealed private parts. Since typical AI deep models take square-shaped images as inputs in terms of pixel counts (often 224×224×3 pixels), we added extra blank pixels in the periphery of many images to avoid excessive stretching of the actual skin lesions during image resizing. Finally, we cropped and resized all the images to 224×224×3 pixels using bilinear interpolation.

#### 4) Augmentation

We performed 19 augmentation operations on the web-scrapped images using Python Imaging Library (PIL) version 9.2.0, and scikit-image library version 0.19.3. to increase the number of images and introduce variability in the data. Our augmentation operations include (1) brightness modification with a randomly generated factor (range [0.5, 2]), (2) color modification with a randomly generated factor (range [0.5, 1.5]), (3) sharpness modification with a randomly generated factor (range [0.5, 2]), (4) image translation along height and width with a randomly generated distance between −25 and 25 pixels, (5) image shearing along height and width with randomly generated parameters, (6) adding Gaussian noise of zero mean and randomly generated variance (range [0.005, 0.02]) to images, (7) adding zero-mean speckle noise and randomly generated variance (range [0.005, 0.02]) to images, (8) adding salt noise to randomly generated number of image pixels (range [2%, 8%]), (9) adding pepper noise to the randomly generated number of image pixels (range [2%, 8%]), (10) adding salt & pepper noise to randomly generated number of image pixels (range [2%, 8%]), (11) modifying the values of the image pixels based on the local variance, (12) blurring an image with a Gaussian kernel with a randomly generated radius (range [1, 3] pixels), (13) contrast modification with a randomly generated factor (range [1, 1.5]), (14) rotating all images by 90°, (15) rotating images at random angle (range [−45°, 45°]), (16) zooming in an image by about 9%, (17) zooming in an image by about 18%, (18) flipping images along the height, and (19) flipping images along the width. In Table I, we show the number of original and augmented images per class in our database. We also show example images after augmentation in Fig. 4. We used these 19 augmentation operations in different combinations that increased the data by 49×.

**TABLE I.**
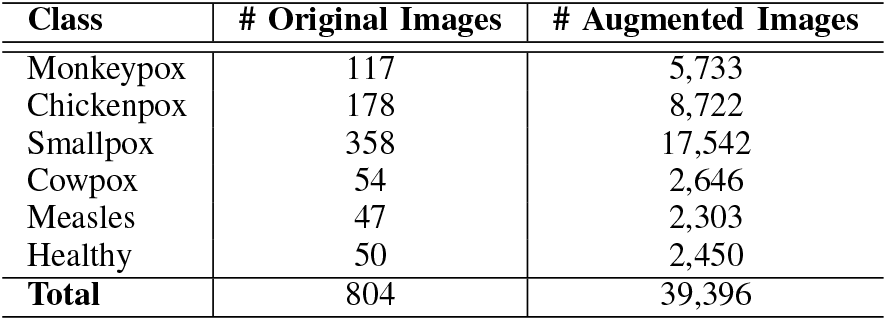
Distribution of image classes in our Monkeypox Skin Image Dataset 2022.

**Fig. 4.**
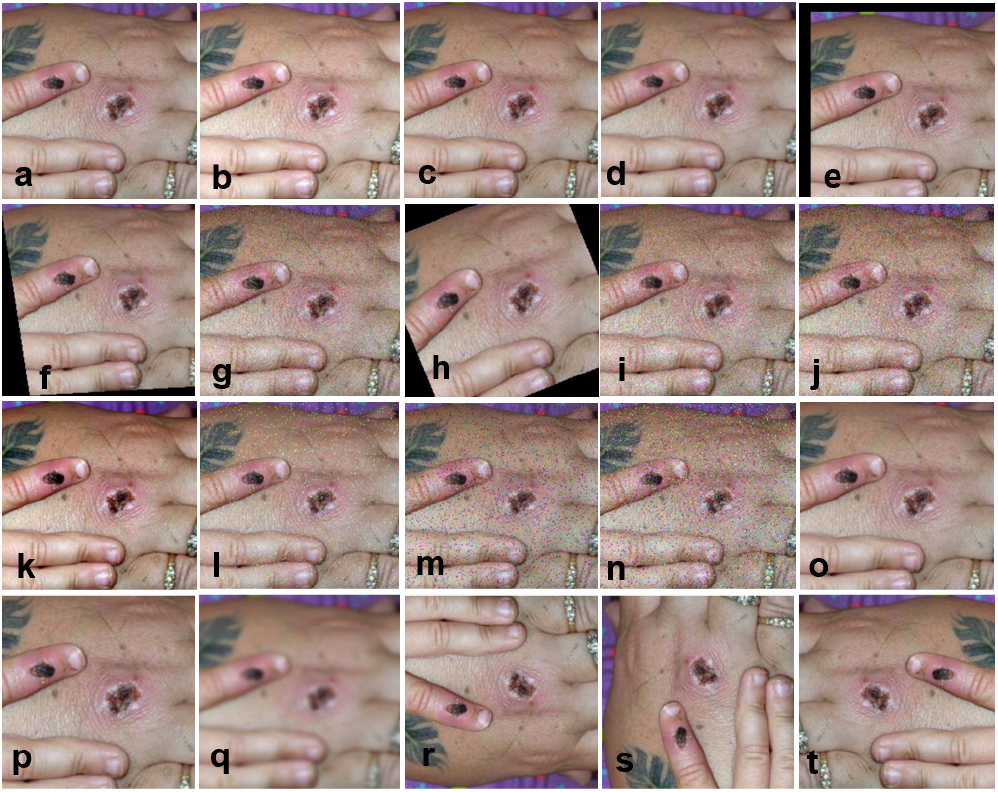
Illustration of (a) an original image and corresponding augmented images by (b) brightness modification, (c) color modification, (d) sharpness modification, (e) translation (f) shear, (g) adding Gaussian noise, (h) random rotation, (i) adding speckle Noise, (j) adding local variance noise, (k) contrast modification (l) adding salt noise, (m) adding pepper noise, (n) adding salt & pepper noise, (o) 9% zooming in, (p) 18% zooming in, (q) Gaussian blurring, (r) flipping along the height, (s) rotation by 90°, and (t) flip along the width.

### B. Experimental Setup

#### 1) Deep Models

We implement seven deep AI models, namely ResNet50 [21], DenseNet121 [22], Inception-V3 [23], SqueezeNet [24], MnasNet-A1 [25], MobileNet-V2 [26], and ShuffleNet-V2-1× [27], to check their feasibility in pox classification task. These models vary in terms of the number of trainable parameters as shown in Table II. Since our dataset is small, we choose deep models of different size ranges (the number of trainable parameters ranges from 1-26 Million) to check the effect of training sample size.

**TABLE II.** Approximate number of trainable parameters in millions (M) of the seven deep AI models used in this study.

| Model | # Parameters (M) |
| --- | --- |
| ResNet50 [21] | 25.6 |
| Inception-V3 [23] | 23.8 |
| DenseNet121 [22] | 7.2 |
| MnasNet-A1 [25] | 3.9 |
| MobileNet-V2 [26] | 3.4 |
| ShuffleNet-V2-1× [27] | 2.3 |
| SqueezeNet [24] | 1.2 |

#### 2) Fine-tuning of Deep Models in Cross-validation

Since our digital skin image dataset is small, we use ImageNet-based pre-trained weights to initialize our deep models and fine-tune on our digital skin image data in 5-fold cross-validation settings. We split our preprocessed original images (shown in Table I second column) in 5 equal folds per class. Since there is a class imbalance among our original data, we use different numbers of augmented images per class during training to make the training data balanced, i.e., all the classes have about the same number of training image samples (~1,700 images per class). We also make sure that augmented images of an original image do not get split into different folds during cross-validation training. In Table III, we show the distribution of the number of images used in validation per class and the number of augmented images used in training per class in each fold.

**TABLE III.** Number of images per class in each fold in a 5-fold cross-validation setup.

| Class | # Validation Images<br>(Validated on) | # Training Images | # Augmented Images<br>(Trained on) |
| --- | --- | --- | --- |
| Monkeypox | 23 | 92 | 1,748 ($=92 \times 19$) |
| Chickenpox | 35 | 140 | 1,680 ($=140 \times 12$) |
| Smallpox | 71 | 284 | 1,704 ($=284 \times 6$) |
| Cowpox | 10 | 40 | 1,760 ($=40 \times 44$) |
| Measles | 9 | 36 | 1,764 ($=36 \times 49$) |
| Healthy | 10 | 40 | 1,760 ($=40 \times 44$) |

### C. Computation Setup

We schedule fine-tuning each deep model for 100 epochs/fold, and the whole 5-fold cross-validation was supposed to take about 1 day/model. We picked the best validation performance in each fold. We chose the Adam optimizer with a learning rate of 0.001. We also chose a batch size of 16 images. We implemented our models in PyTorch version 1.6.0 and Python version 3.8.10. The training was performed on a workstation with an Intel E5-2650 v4 Broadwell 2.2 GHz processor, an Nvidia P100 Pascal GPU with 16 GB of VRAM, and 64 GB of RAM.

## III. Results

In this section, we present the comparative classification performance of our seven state-of-the-art deep AI models as summarized in Table II. In Figs. 5, 6, 7, 8, 9, 10, and 11, we show the confusion matrices for 5-fold cross-validation predictions by ResNet50, Inception-V3, DenseNet121, MnasNet-A1, MobileNet-V2, ShuffleNet-V2-1×, and SqueezeNet, respectively. We see many misclassifications for disease classes in these figures for all the deep models, except for the healthy skin class. The least number of misclassifications for all 5 folds is seen for ShuffleNet-V2 (see Fig. 10).

**Fig. 5.**
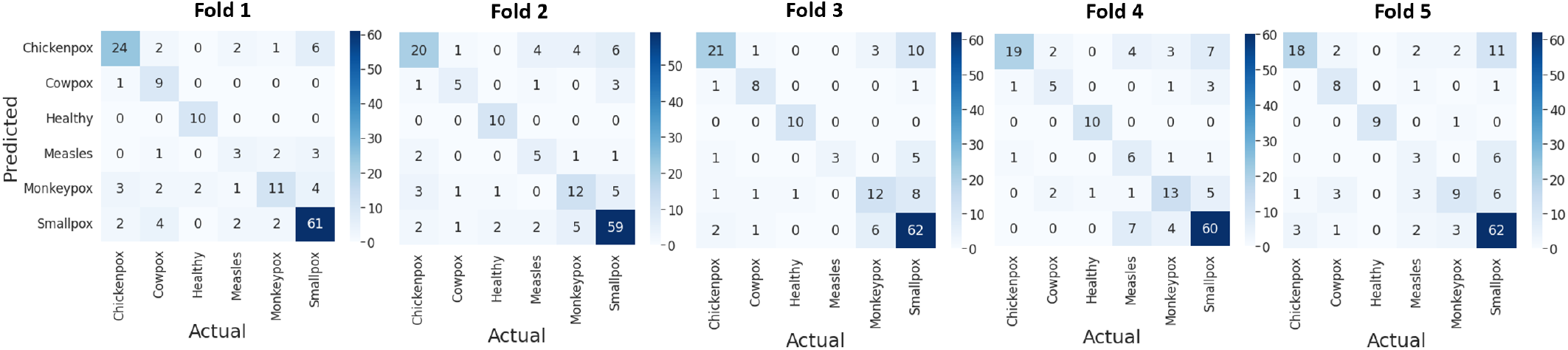
Confusion matrices of prediction by ResNet50 in 5-fold cross-validation.

**Fig. 6.**
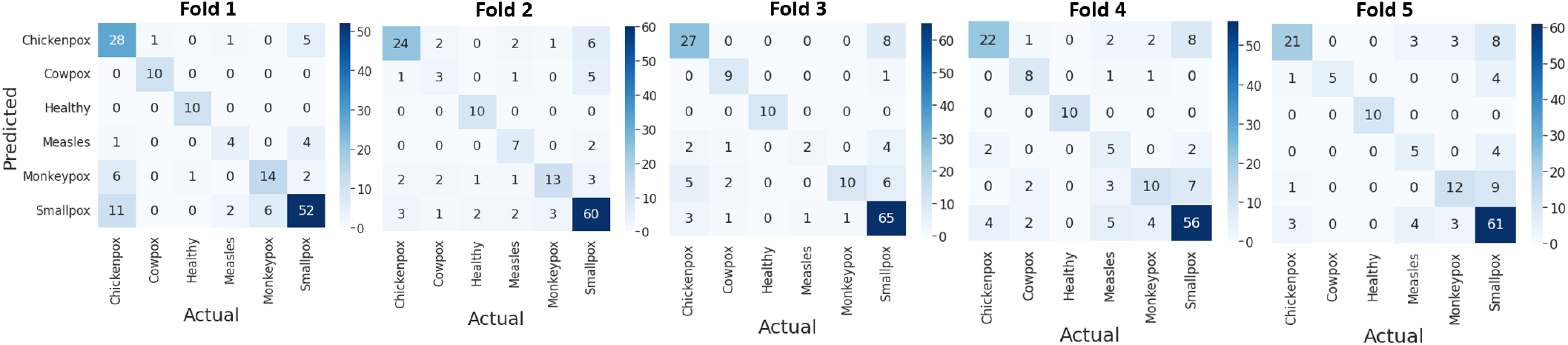
Confusion matrices of prediction by Inception-V3 in 5-fold cross-validation.

**Fig. 7.**
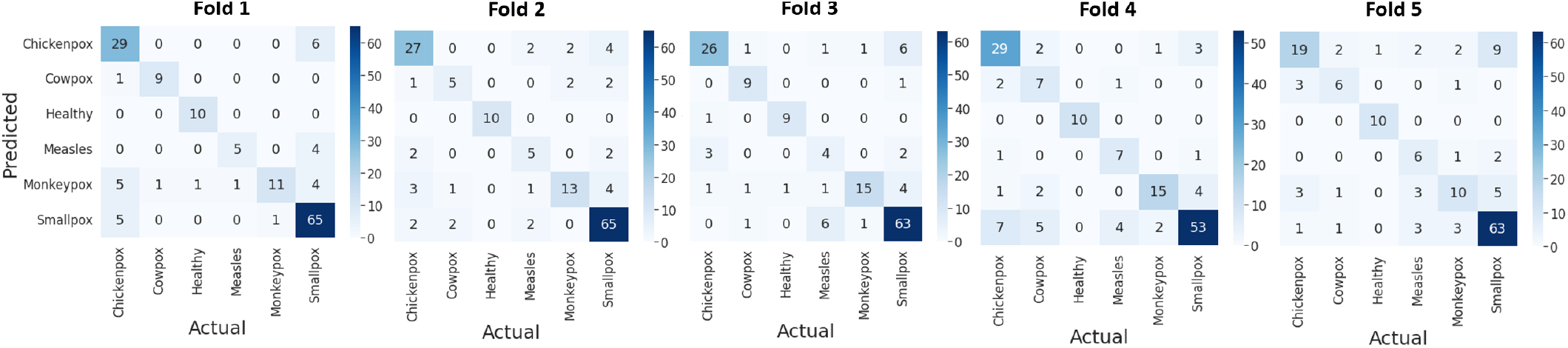
Confusion matrices of prediction by DenseNet121 in 5-fold cross-validation.

**Fig. 8.**
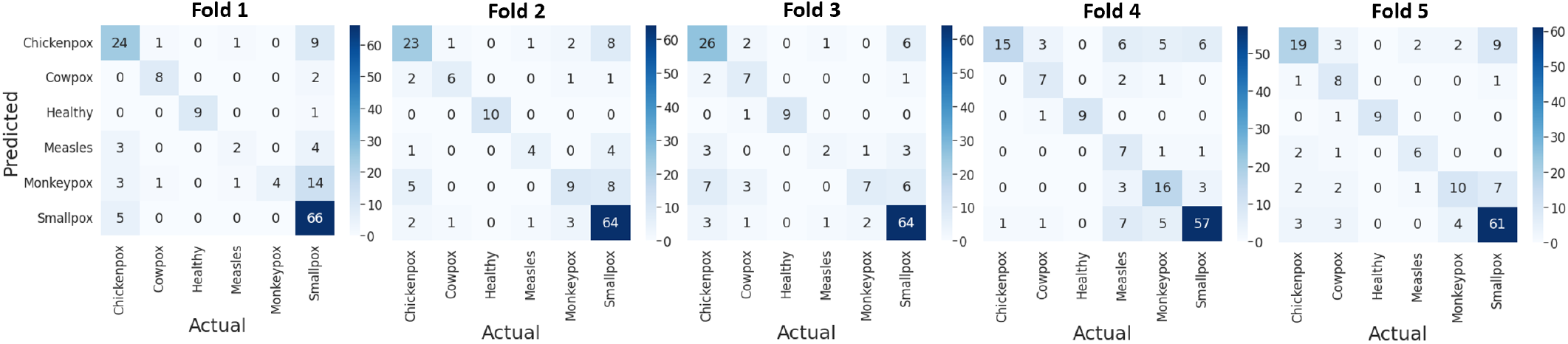
Confusion matrices of prediction by MnasNet-A1 in 5-fold cross-validation.

**Fig. 9.**
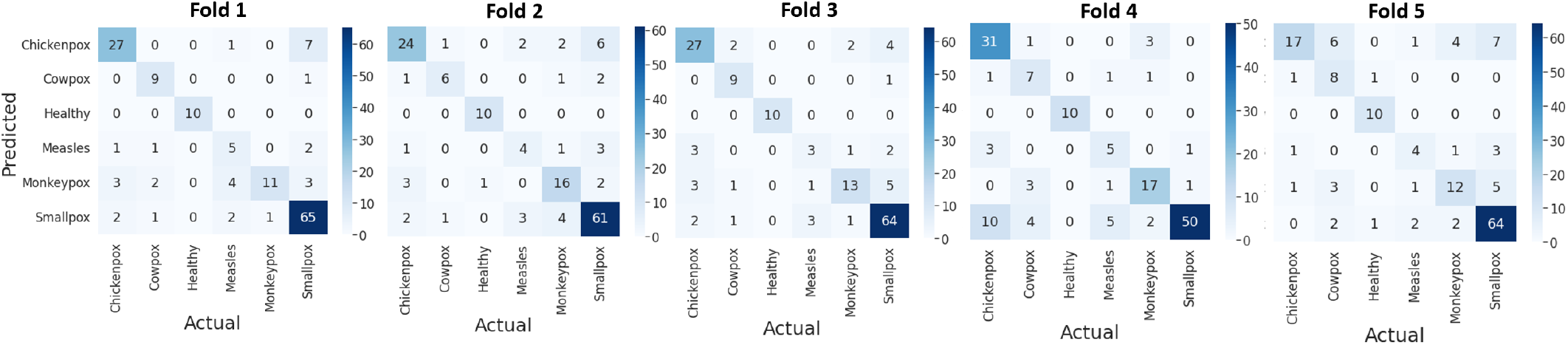
Confusion matrices of prediction by MobileNet-V2 in 5-fold cross-validation.

**Fig. 10.**
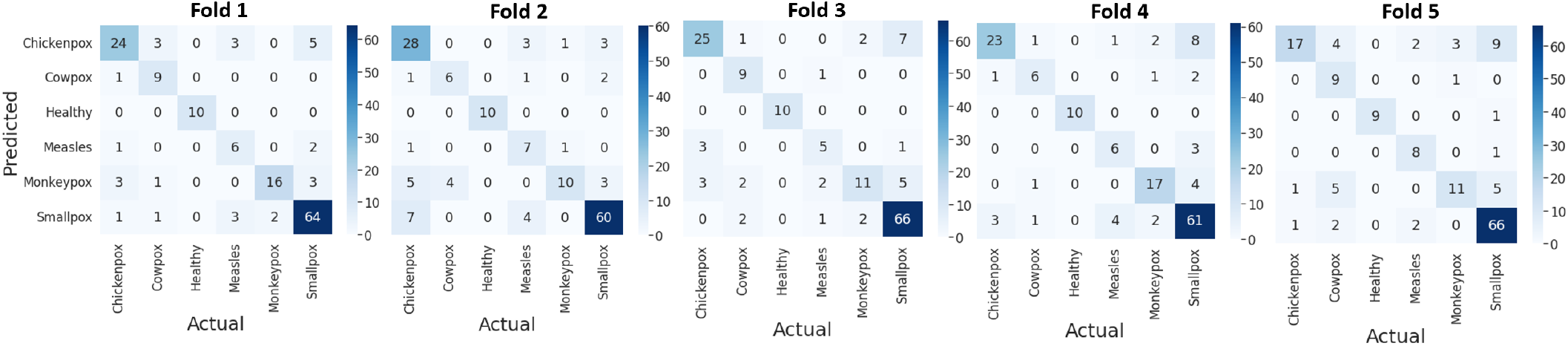
Confusion matrices of prediction by ShuffleNet-V2-1× in 5-fold cross-validation.

**Fig. 11.**
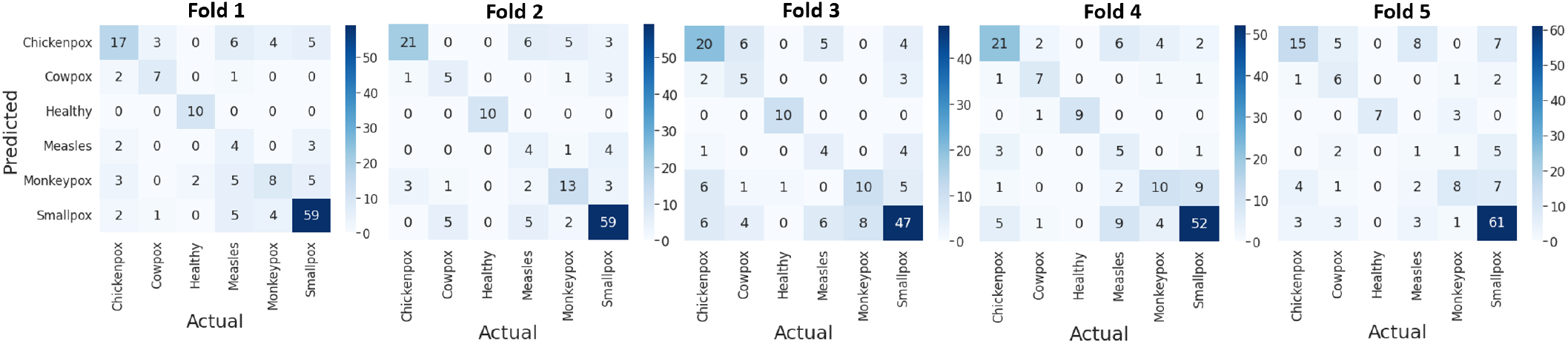
Confusion matrices of prediction by SqueezeNet in 5-fold cross-validation.

We also present the quantitative comparison of mean precision, mean recall, mean F1 score and mean accuracy, estimated over the 5-fold cross-validation, for all classes by all deep AI models in Table IV. Here, we see that the best accuracy is achieved by the ShuffleNet-V2 (79%). ShuffleNet-V2 has fewer trainable parameters (i.e., 2.3 M) than ResNet50 (25.6 M), Inception-V3 (23.8 M), DenseNet121 (7.2 M), MnasNet-A1 (3.9 M) and MobileNet-V2 (3.4 M) (see Table II). From our observation of the prediction performance by different deep models, as summarized in Table IV, we hypothesize that models with a larger number of trainable parameters may be under fitted due to the small training sample size. On the other hand, although the SqueezeNet has fewer trainable parameters (i.e., 1.2 M) than the ShuffleNet-V2 (2.3 M), it is overfitted on the small number of training samples, thus resulting in worse prediction performance on the validation data.

**TABLE IV.** Quantitative comparison of mean precision, mean recall, mean F1 score and mean accuracy over the 5-fold cross-validation.

| Methods | Class | Mean Precision | Mean Recall | Mean F1 Score | Mean Accuracy |
| --- | --- | --- | --- | --- | --- |
| ResNet50 | Chickenpox | 0.80 | 0.58 | 0.68 | 0.72 |
|  | Cowpox | 0.58 | 0.70 | 0.64 |  |
|  | Healthy | 0.88 | 0.98 | 0.92 |  |
|  | Measles | 0.38 | 0.44 | 0.41 |  |
|  | Monkeypox | 0.59 | 0.51 | 0.55 |  |
|  | Smallpox | 0.77 | 0.86 | 0.81 |  |
| Inception-V3 | Chickenpox | 0.73 | 0.70 | 0.71 | 0.71 |
|  | Cowpox | 0.70 | 0.70 | 0.70 |  |
|  | Healthy | 0.93 | 1.00 | 0.96 |  |
|  | Measles | 0.45 | 0.51 | 0.48 |  |
|  | Monkeypox | 0.71 | 0.53 | 0.61 |  |
|  | Smallpox | 0.77 | 0.83 | 0.80 |  |
| DenseNet121 | Chickenpox | 0.76 | 0.74 | 0.75 | 0.78 |
|  | Cowpox | 0.64 | 0.72 | 0.68 |  |
|  | Healthy | 0.94 | 0.98 | 0.96 |  |
|  | Measles | 0.50 | 0.60 | 0.55 |  |
|  | Monkeypox | 0.71 | 0.53 | 0.61 |  |
|  | Smallpox | 0.83 | 0.87 | 0.85 |  |
| MnasNet-A1 | Chickenpox | 0.70 | 0.61 | 0.65 | 0.72 |
|  | Cowpox | 0.58 | 0.72 | 0.64 |  |
|  | Healthy | 1.00 | 0.92 | 0.96 |  |
|  | Measles | 0.44 | 0.47 | 0.45 |  |
|  | Monkeypox | 0.63 | 0.41 | 0.50 |  |
|  | Smallpox | 0.77 | 0.88 | 0.82 |  |
| MobileNet-V2 | Chickenpox | 0.77 | 0.72 | 0.74 | 0.77 |
|  | Cowpox | 0.57 | 0.78 | 0.66 |  |
|  | Healthy | 0.94 | 1.00 | 0.97 |  |
|  | Measles | 0.44 | 0.47 | 0.45 |  |
|  | Monkeypox | 0.73 | 0.62 | 0.67 |  |
|  | Smallpox | 0.85 | 0.86 | 0.85 |  |
| ShuffleNet-V2 | Chickenpox | 0.79 | 0.67 | 0.72 | 0.79 |
|  | Cowpox | 0.58 | 0.78 | 0.67 |  |
|  | Healthy | 1.00 | 0.98 | 0.99 |  |
|  | Measles | 0.54 | 0.71 | 0.62 |  |
|  | Monkeypox | 0.79 | 0.58 | 0.67 |  |
|  | Smallpox | 0.83 | 0.89 | 0.86 |  |
| SqueezeNet | Chickenpox | 0.67 | 0.54 | 0.60 | 0.65 |
|  | Cowpox | 0.45 | 0.60 | 0.52 |  |
|  | Healthy | 0.94 | 0.92 | 0.93 |  |
|  | Measles | 0.20 | 0.40 | 0.27 |  |
|  | Monkeypox | 0.55 | 0.44 | 0.49 |  |
|  | Smallpox | 0.79 | 0.78 | 0.78 |  |

We also present classification performance by an ensemble of our implemented and fine-tuned seven deep models, where we use majority voting to make the prediction decision. In Fig. 12, we show the confusion matrices for 5-fold cross-validation predictions by the ensemble of ResNet50, Inception-V3, DenseNet121, MnasNet-A1, MobileNet-V2, ShuffleNet-V2-1× and SqueezeNet. We see in these confusion matrices that the ensemble approach reduces the missclassifications significantly, compared to those seen in Figs. 5, 6, 7, 8, 9, 10 and 11 for individual deep models. Furthermore, we present the quantitative comparison of mean precision, mean recall, mean F1 score, and mean accuracy, estimated over the 5-fold cross-validation, for all classes using the ensemble approach in Table V. We see in this table that the ensemble approach shows the best performance in terms of all the metrics, notably in terms of accuracy (83%), compared to those by individual deep models as summarized in Table IV.

**Fig. 12.**
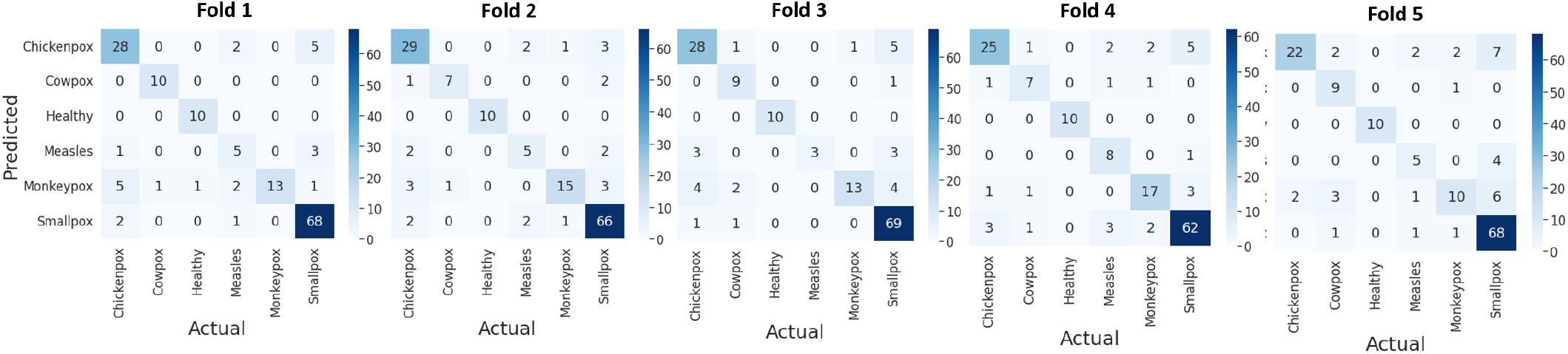
Confusion matrices of prediction by majority voting in 5-fold cross-validation.

**TABLE V.** Quantitative performance by the ensemble method in terms of mean precision, mean recall, mean F1 score, and mean accuracy over the 5-fold cross-validation.

| Methods | Class | Mean Precision | Mean Recall | Mean F1 Score | Mean Accuracy |
| --- | --- | --- | --- | --- | --- |
| Majority Voting | Chickenpox | 0.81 | 0.75 | 0.78 | 0.83 |
|  | Cowpox | 0.74 | 0.84 | 0.79 |  |
|  | Healthy | 0.98 | 1.00 | 0.99 |  |
|  | Measles | 0.58 | 0.58 | 0.58 |  |
|  | Monkeypox | 0.85 | 0.61 | 0.71 |  |
|  | Smallpox | 0.85 | 0.94 | 0.89 |  |

## IV. Conclusion

In this paper, we tested the feasibility of using seven state-of-the-art AI deep models in classifying different pox types and Measles from the digital skin images of lesions and rashes. We built and utilized a digital skin database containing skin lesion/rash images of five different diseases, namely, Monkeypox, Chickenpox, Smallpox, Cowpox, and Measles. We also chose seven state-of-the-art deep models that vary in terms of the number of trainable parameters, ranging from 1 to 26 million. Our 5-fold cross-validation experiments showed that AI deep models have the ability to distinguish among different pox types using digital skin images of pox/Measles lesions and rashes. We also observed that deep models tend to overfit or underfit, perhaps due to the trade-off between the training sample size and the number of trainable parameters in a model. Thus, to achieve better classification accuracy and harness the full strength of state-of-the-art deep models, we must ensure a larger sample size for model training. We also observed that lighter deep models, having fewer trainable parameters, also have potential in pox classification, which could be used on smartphones for Monkeypox diagnosis in the latest outbreak. Furthermore, Monkeypox detection based on digital skin images can facilitate remote diagnosis by healthcare professionals, leading to early isolation of patients and effective containment of community spread.

